# SARS-CoV-2 escape *in vitro* from a highly neutralizing COVID-19 convalescent plasma

**DOI:** 10.1101/2020.12.28.424451

**Authors:** Emanuele Andreano, Giulia Piccini, Danilo Licastro, Lorenzo Casalino, Nicole V. Johnson, Ida Paciello, Simeone Dal Monego, Elisa Pantano, Noemi Manganaro, Alessandro Manenti, Rachele Manna, Elisa Casa, Inesa Hyseni, Linda Benincasa, Emanuele Montomoli, Rommie E. Amaro, Jason S. McLellan, Rino Rappuoli

## Abstract

To investigate the evolution of SARS-CoV-2 in the immune population, we co-incubated authentic virus with a highly neutralizing plasma from a COVID-19 convalescent patient. The plasma fully neutralized the virus for 7 passages, but after 45 days, the deletion of F140 in the spike N-terminal domain (NTD) N3 loop led to partial breakthrough. At day 73, an E484K substitution in the receptor-binding domain (RBD) occurred, followed at day 80 by an insertion in the NTD N5 loop containing a new glycan sequon, which generated a variant completely resistant to plasma neutralization. Computational modeling predicts that the deletion and insertion in loops N3 and N5 prevent binding of neutralizing antibodies. The recent emergence in the United Kingdom and South Africa of natural variants with similar changes suggests that SARS-CoV-2 has the potential to escape an effective immune response and that vaccines and antibodies able to control emerging variants should be developed.

**One Sentence Summary:** Three mutations allowed SARS-CoV-2 to evade the polyclonal antibody response of a highly neutralizing COVID-19 convalescent plasma.

---

The SARS-CoV-2 virus, causative agent of COVID-19, accounts for over 78.5 million cases of infections and almost 2 million deaths worldwide. Thanks to an incredible scientific and financial effort, several prophylactic and therapeutic tools, such as vaccines and monoclonal antibodies (mAbs), have been developed in less than one year to combat this pandemic (*1–4*). The main target of vaccines and mAbs is the SARS-CoV-2 spike protein (S-protein), a large class I trimeric fusion protein which plays a key role in viral pathogenesis (*3, 5, 6*). The SARS-CoV-2 S-protein is composed of two subunits: S1, which contains the receptor-binding domain (RBD) responsible for the interaction with receptors on the host cells, and S2, which mediates membrane fusion and viral entry (*7, 8*). The S1 subunit presents two highly immunogenic domains, the N-terminal domain (NTD) and the RBD, which are the major targets of polyclonal and monoclonal neutralizing antibodies (*4, 9, 10*). The continued spread in immune-competent populations has led to adaptations of the virus to the host and generation of new SARS-CoV-2 variants. Indeed, S-protein variants have been recently described in the United Kingdom and South Africa (*11, 12*), and the Global Initiative on Sharing All Influenza Data (GISAID) database, reports more than 1,100 amino acid changes in the S-protein (*13, 14*).

An important question for vaccine development is whether the authentic virus, under the selective pressure of the polyclonal immune response in convalescent or vaccinated people, can evolve to escape herd immunity and antibody treatment. To address this question, we collected plasma from 20 convalescent patients and incubated the authentic SARS-CoV-2 wild-type (WT) virus for more than 90 days in the presence of a potent neutralizing plasma. Enzyme-linked immunosorbent assay (ELISA) showed that all plasmas collected bound the SARS-CoV-2 S-protein trimer and most of them also bound the S1 and S2 subunits. However, a broad range of reactivity profiles were noticed ranging from weak binders with titers of 1/80 to strong binders with titers of 1/10240 (**Fig S1A**; **Table S1**). PT008, PT009, PT015, PT122 and PT188 showed the strongest binding towards the S trimer and among them PT188 had also the highest binding to the S1 and S2 subunits. All but one plasma sample (PT103) were able to bind the S-protein S1 subunit. Neutralization activity tested against the SARS-CoV-2 WT and D614G variant also showed variable titers. Most of the plasma samples neutralized the viruses with titers ranging from 1/20 to 1/320. Four samples had extremely low titers (1/10), whereas sample PT188 showed extremely high titers (1/10240). Four plasma samples did not show neutralization activity against the SARS-CoV-2 WT and SARS-CoV-2 D614G variant. Plasma from subject PT188, which had the highest neutralizing titer and ELISA binding reactivity (**Fig. S1B, C, D**; **Table 1**), was selected to test whether SARS-CoV-2 can evolve to escape a potent humoral immunity.

Two-fold dilutions of plasma PT188 ranging from 1/10 to 1/20480 were co-incubated with 10^5^ TCID_50_ of the wild type virus in a 24-well plate. This viral concentration is approximately three logs more than what is conventionally used in microneutralization assays (*15–19*). The plasma/virus mixture was co-incubated for 5–8 days. Then, the first well showing cytopathic effect (CPE) was diluted 1:100 and incubated again with serial dilutions of plasma PT188 (**Fig. 1A; Table S2**). For 6 passages and 38 days PT188 plasma neutralized the virus with a titer of 1/640 and did not show any sign of escape. However, after 7 passages and 45 days, the neutralizing titer decreased to 1/320. Sequence analyses revealed a deletion of the phenylalanine in position 140 (F140) on the S-protein NTD N3 loop in 36% of the virions (**Fig. 1, B and C; Table S2**). In the subsequent passage (P8), this mutation was observed in 100% of the sequenced virions and an additional 2-fold decrease in neutralization activity was observed reaching an overall neutralization titer of 1/160. Following this initial breakthrough, a second mutation occurred after 12 passages and 80 days of plasma/virus co-incubation (P12). This time, the glutamic acid in position 484 of the RBD was substituted with a lysine (E484K). This mutation occurred in 100% of sequenced virions and led to a 4-fold decrease in neutralization activity which reached a titer of 1/40 (**Fig. 1, B and C; Table S2**). The E484K substitution was rapidly followed by a third and final change comprising an 11-amino-acid insertion between Y248 and L249 in the NTD N5 loop (_248a_KTRNKSTSRRE_248k_). The insertion contained an N-linked glycan sequon (_248d_NKS_248f_), and this viral variant resulted in complete abrogation of neutralization activity by the PT188 plasma sample. Initially this insertion was observed in only 49% of the virions but when the virus was kept in culture for another passage (P14) the insertion was fully acquired by the virus (**Fig. 1, B and C; Table S2**).

**Fig. 1.**
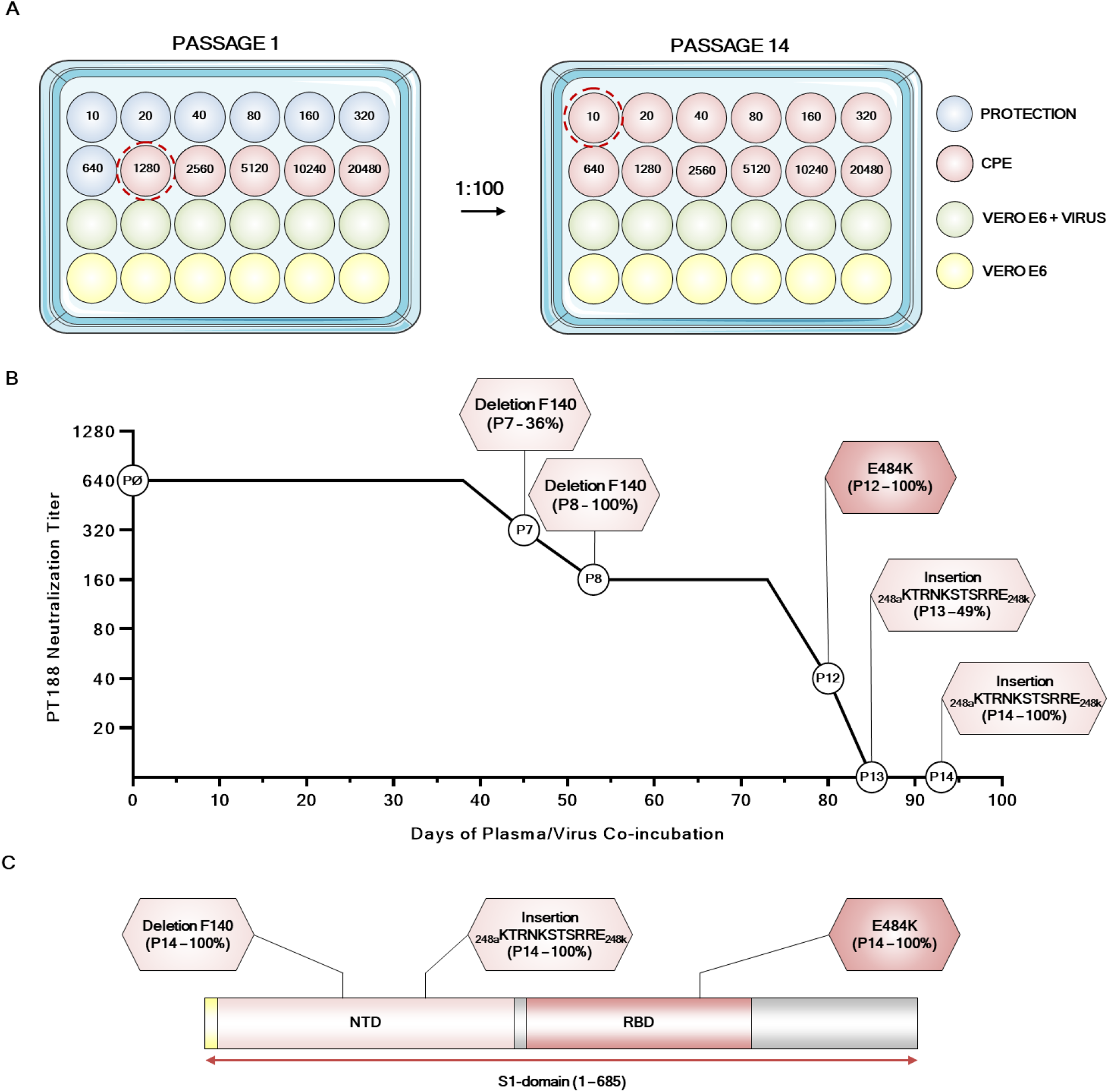
Evolution of an authentic SARS-CoV-2 escape mutant. (A) Schematic representation of the 24-well plate format used to select the authentic SARS-CoV-2 escape mutant. Blue, red, green and yellow wells show feeder cells protect from PT188 neutralization, CPE, authentic virus on Vero E6 cells and Vero E6 alone, respectively. (B) The graph shows the PT188 neutralization titer after each mutation acquired by the authentic virus. Specific mutations, fold decrease and days to which the mutations occur are reported in the figure. (C) SARS-CoV-2 S-protein gene showing type, position of mutations and frequency of mutations.

To evaluate the ability of the SARS-CoV-2 PT188 escape mutant (PT188-EM) to evade the polyclonal antibody response, all twenty plasma samples from COVID-19 convalescent patients were tested in a traditional CPE-based neutralization assay against this viral variant using the virus at 100 TCID_50_. All samples showed at least a 2-fold decrease in neutralization activity against SARS-CoV-2 PT188-EM (**Fig. 2A; Fig. S1B, C, D; Table S1**). As expected, the plasma used to select the escape mutant showed the biggest neutralization decrease against this escape mutant with a 256-fold decrease compared to wild-type SARS-CoV-2. Plasma PT042, PT006, PT005, PT012 and PT041 also showed a substantial drop in neutralization efficacy (**Table S1**). In addition, we observed that a higher response towards the S-protein S1-subunit correlates with loss of neutralization activity against SARS-CoV-2 PT188-EM (**Fig. S2A**) whereas a high response towards the S-protein S2-subunit did not show correlation (**Fig. S2B**).

**Fig. 2.**
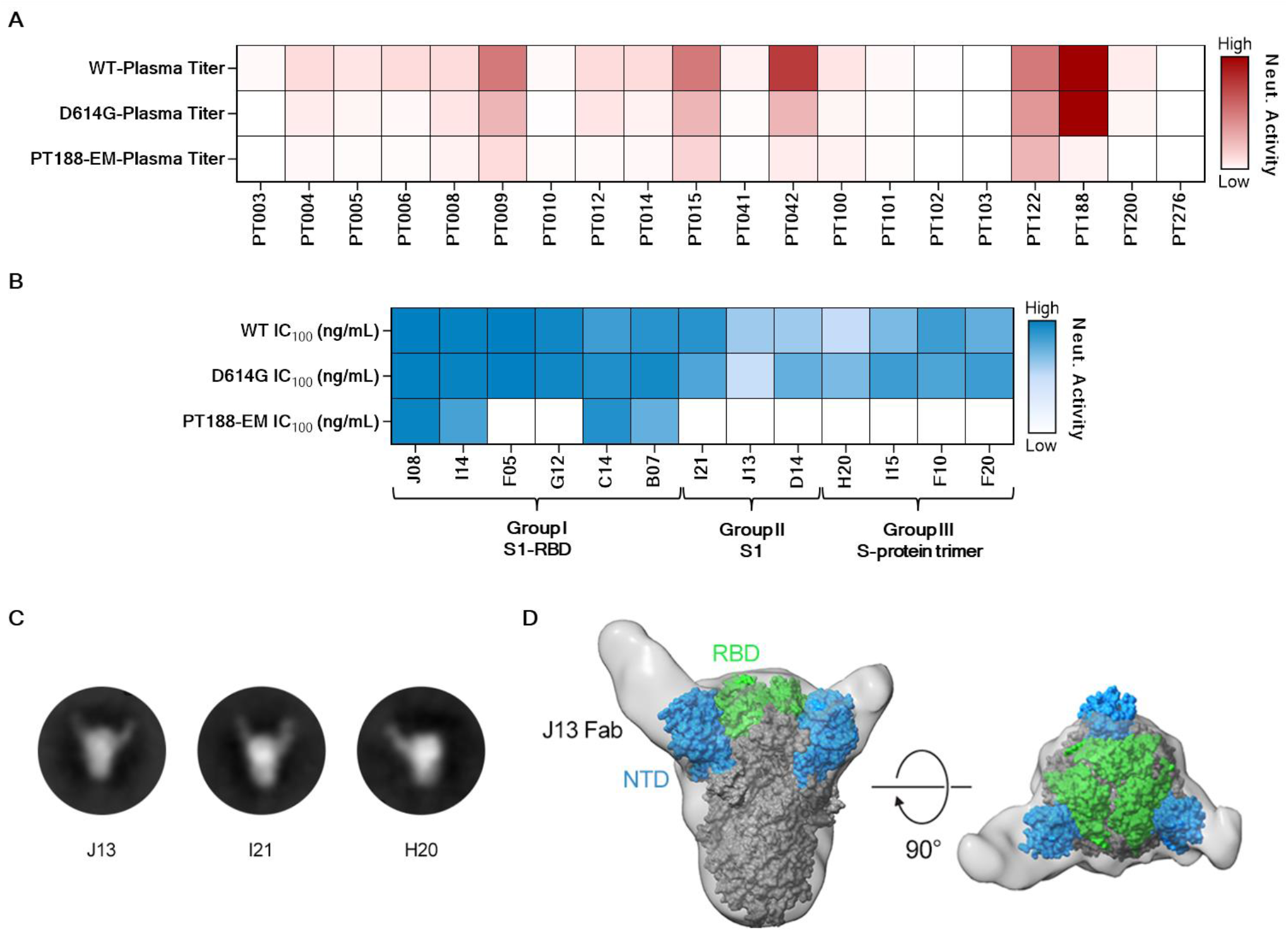
Neutralization efficacy of plasma and thirteen mAbs to SARS-CoV-2 PT188-EM. (A) Heat-map showing the neutralization activity of tested plasma samples to the SARS-CoV-2 WT, D614G and PT188-EM variants. (B) Heat-maps showing neutralization profiles of tested mAbs. (C) Negative stain EM 2D class averages showing J13, I21 and H20 Fabs bound to the SARS-CoV-2 S-protein. (D) 3D reconstruction of J13 bound to the NTD domain of the S-protein viewed looking along (left panel) or toward (right panel) the viral membrane.

We also tested a previously identified panel of thirteen neutralizing monoclonal antibodies (nAbs) (*19*) by CPE-based neutralization assay to assess their neutralization efficacy against SARS-CoV-2 PT188-EM. These antibodies were classified in three groups based on their binding profiles to the S-protein. Group I nAbs were able to bind the S1-RBD, Group II targeted the S1-subunit but not the RBD, and Group III nAbs were specific for the S-protein trimer **(Table S3**). These antibodies also showed a variable neutralization potency against the SARS-CoV-2 WT and D614G viruses ranging from 3.9 ng/mL to 500.0 ng/mL (**Fig. 2B; Fig. S2E, F, G; Table S3**). The three mutations selected by SARS-CoV-2 PT188-EM to escape the highly neutralizing plasma completely abrogated the neutralization activity of two of the six tested RBD-directed antibodies (F05 and G12) (**Fig. 2B; Fig. S2E, F, G; Table S3**), suggesting that their epitopes include E484. In contrast, the extremely potent neutralizing antibody J08 was the most potently neutralizing antibody against this escape mutant with an IC_100_ of 22.1 ng/mL. Interestingly, the S1-RBD-directed antibody C14 showed a 2-fold increase in neutralization activity compared to the SARS-CoV-2 WT virus whereas I14 and B07 showed a 16- and 2-fold decrease, respectively. All tested antibodies derived from Group II (S1-specific not RBD) and Group III (S-protein trimer specific) completely lost their neutralization ability against SARS-CoV-2 PT188-EM (**Fig. 2B; Fig. S2E, F, G; Table S3**). To better understand the abrogation of activity of some of the tested antibodies, J13, I21 and H20 were co-complexed with SARS-CoV-2 WT S-protein and structurally evaluated by negative-stain EM. 2D class averages of the three tested antibodies showed that they all bind to the NTD of the S-protein (**Fig. 2C**). A 3D reconstruction for the J13 Fab complex provided further evidence that this antibody binds to the NTD (**Fig. 2D**).

Computational modeling and simulation of the WT and PT188-EM spikes provides a putative structural basis for understanding antibody escape. The highly antigenic NTD is more extensively mutated, containing both the F140 deletion as well as the 11-amino-acid insertion in loop N5 that introduces a novel N-glycan sequon at position N248d (**Fig. 3, A - C)**. In contrast, the single mutation in the RBD (E484K) swaps the charge of the sidechain, which would significantly alter the electrostatic complementarity of antibody binding to this region (**Fig. 3D**). Upon inspection of molecular dynamics simulations of the NTD escape mutant model, we hypothesize that the F140 deletion alters the packing of the N1, N3 and N5 loops (**Fig. S3**), where the loss of the bulky aromatic sidechain would overall reduce the stability of this region (**Table S1**). Subsequently, the extensive insertion within the N5 loop appears to remodel this critical antigenic region, predicting substantial steric occlusion with antibodies targeting this epitope, such as antibody 4A8 (**Fig. 3B**) (*20*). Furthermore, introduction of a new N-glycan at position N248d (mutant numbering scheme) would effectively eliminate neutralization by such antibodies (**Fig. 3B and S4**).

**Fig. 3.**
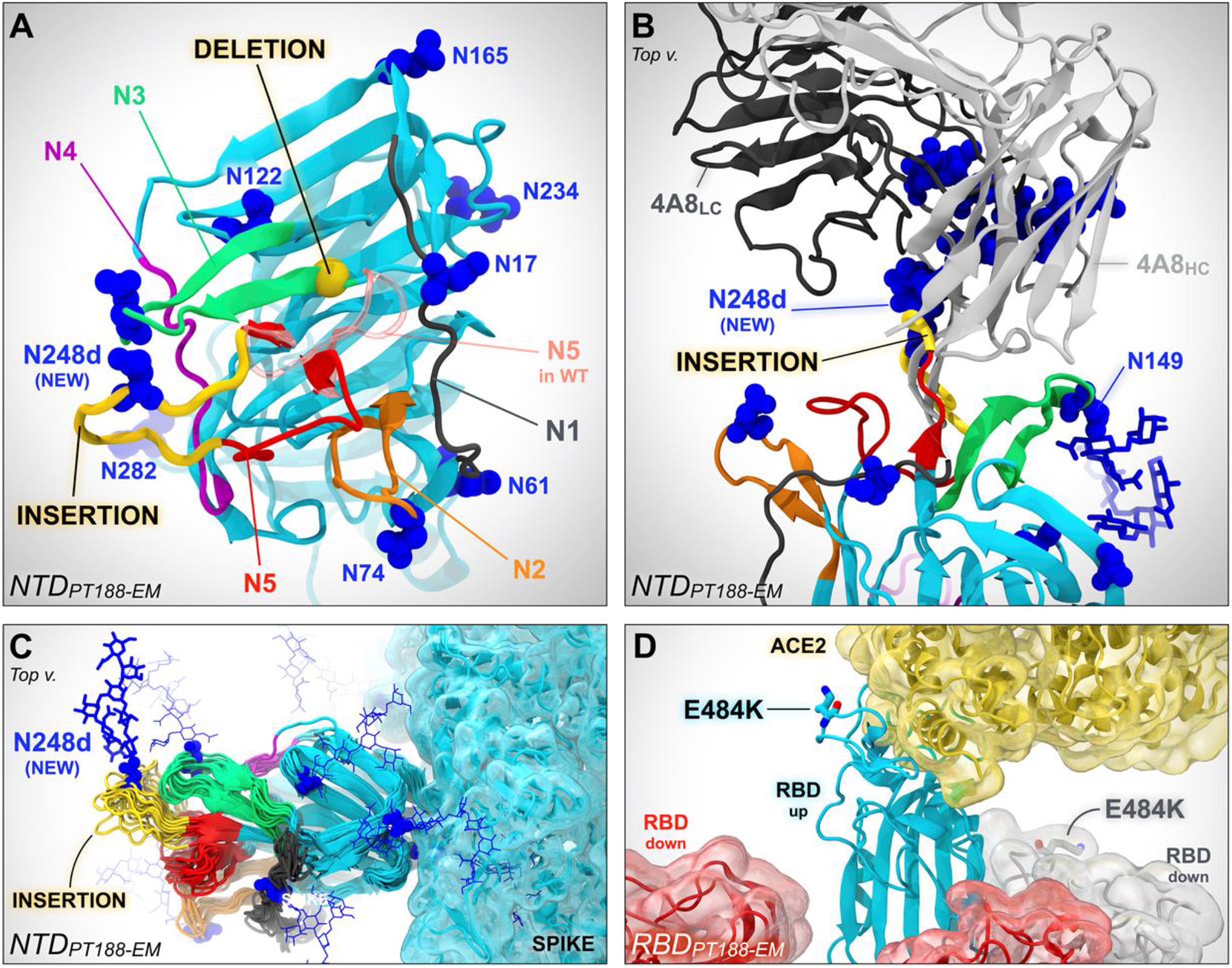
*In-silico* modeling of the PT188-EM spike NTD and RBD. (A) *In-silico* model of the NTD of the SARS-CoV-2 PT188-EM spike protein based on PDB id 7JJI. This model accounts for the 11-amino-acid insertion (yellow ribbon) and F140 deletion (highlighted with a yellow bead). N5 loop as in the wild type cryo-EM structure (PDB id: 7JJI) is shown as a transparent red ribbon. (B) Close-up of the PT188-EM spike NTD model in complex with antibody 4A8. Both heavy chain (HC, light gray) and light chain (LC, dark gray) of 4A8 are shown. The 11-amino-acid insertion (yellow ribbon) within N5 loop introduces a new N-linked glycan (N248d) that sterically clashes with 4A8, therefore disrupting the binding interface. The N-glycan at position N149 is however compatible with 4A8 binding. (C) Conformational dynamics of the PT188-EM spike NTD model resulting from 100 ns of molecular dynamics simulation is shown by overlaying multiple frames along the generated trajectory. (D) *In-silico* model of the PT188-EM spike RBD based on PDB id 6M17, where the E484K mutation is shown with licorice representation.

To determine the extent to which the escape mutations were detrimental to the infectivity of SARS-CoV-2 PT188-EM, the viral fitness was evaluated. Four different measures were assessed: visible CPE, viral titer, RNA-dependent RNA polymerase (RdRp) and nucleocapsid (N) RNA detection by reverse transcription polymerase chain reaction (RT-PCR) (**Fig. S5**). Initially, the SARS-CoV-2 WT virus and the PT188-EM variant were inoculated at a multiplicity of infection (MOI) of 0.001 on Vero E6 cells. Every day, for four consecutive days, a titration plate was prepared and optically assessed after 72 hours of incubation to evaluate the CPE effect on Vero E6 cells and viral titer. Furthermore, the RNA was extracted to asses RdRp and N-gene levels in the supernatant. We collected pictures at 72 hours post-infection to evaluate the morphological status of non-infected Vero E6 cells and the CPE on infected feeder cells. Vero E6 cells were confluent at 72 hours and no sign of CPE was optically detectable (**Fig S5A**). On the contrary, SARS-CoV-2 WT and PT188-EM showed significant and comparable amount of CPE (**Fig. S5A**). Viral titers were evaluated for both SARS-CoV-2 WT and PT188-EM, and no significant differences were observed as the viruses showed almost identical growth curves (**Fig. S5B**). A similar trend was observed when RdRp and N-gene levels in the supernatant were detected, even if slightly higher levels of RdRp and N-gene were detectable for SARS-CoV-2 PT188-EM at Day 0 and Day 1 (**Fig. S5C**). Finally, strong correlations between viral titers and RdRp/N-gene levels were observed for both SARS-CoV-2 WT and PT188-EM (**Fig. S5D - E**).

In conclusion, we have shown that the authentic SARS-CoV-2 virus, if constantly pressured, has the ability to escape even a potent polyclonal serum targeting multiple neutralizing epitopes. These results are remarkable because while escape mutants can be easily isolated when viruses are incubated with single monoclonal antibodies, usually a combination of two mAbs is sufficient to eliminate the evolution of escape variants and because SARS-CoV-2 shows a very low estimated evolutionary rate of mutation as this virus encodes a proofreading exoribonuclease machinery (*21, 22*)(*23, 24*). The recent isolation of SARS-CoV-2 variants in the United Kingdom and South Africa with deletions in or near the NTD loops shows that what we describe here can occur in the human population. The ability of the virus to adapt to the host immune system was also observed in clinical settings where an immunocompromised COVID-19 patient, after 154 days of infection, presented different variants of the virus including the E484K substitution (*25*). Therefore, we should be prepared to deal with virus variants that may be selected by the immunity acquired from infection or vaccination. This can be achieved by developing second-generation vaccines and monoclonal antibodies, possibly targeting universal epitopes and able to neutralize emerging variants of the virus.

Our data also confirm that the SARS-CoV-2 neutralizing antibodies acquired during infection target almost entirely the NTD and the RBD. In the RBD, the possibility to escape is limited and the mutation E484K that we found is one of the most frequent mutations to escape monoclonal antibodies and among the most common RBD mutations described in experimental settings as well as in natural isolates posted in the GISAD database (*13, 26, 27*). This is likely due to residue E484 being targeted by antibodies derived from IGHV3-53 and closely related IGHV3-66 genes, which are the most common germlines for antibodies directed against the RBD (*28*). On the other hand, the NTD loops can accommodate many different changes, such as insertions, deletions and amino-acid alterations. Interestingly, in our case, the final mutation contained an insertion carrying an N-glycosylation site which has the potential to hide or obstruct the binding to neutralizing epitopes. The introduction of a glycan is a well-known immunogenic escape strategy described in influenza (*29*), HIV-1 and other viruses (*30–32*), although to our knowledge this finding presents the first patient-derived escape mutant utilizing this mechanism for SARS-CoV-2. Surprisingly, only three mutations, which led to complete rearrangement of NTD N3 and N5 loops and substitution to a key residue on the RBD, were sufficient to eliminate the neutralization ability of a potent polyclonal serum. Fortunately, not all plasma and mAbs tested were equally affected by the three mutations suggesting that natural immunity to infection can target additional epitopes that can still neutralize the PT188-EM variant. Therefore, it will be important to closely monitor which epitopes on the S-protein are targeted by the vaccines against SARS-CoV-2 that are going to be deployed in hundreds of millions of people around the world.

## ACKNOWLEDGMENTS

This work was funded by the European Research Council (ERC) advanced grant agreement number 787552 (vAMRes). This publication was supported by funds from the “Centro Regionale Medicina di Precisione” and by all the people who answered the call to fight with us the battle against SARS-CoV-2 with their kind donations on the platform ForFunding (https://www.forfunding.intesasanpaolo.com/DonationPlatform-ISP/nav/progetto/id/3380).

This publication was supported by the European Virus Archive goes Global (EVAg) project, which has received funding from the European Union’s Horizon 2020 research and innovation programme under grant agreement No 653316. This publication was supported by the COVID-2020-12371817 project, which has received funding from the Italian Ministry of Health. This work was supported by NIH GM132826, NSF RAPID MCB-2032054, an award from the RCSA Research Corp., and a UC San Diego Moore’s Cancer Center 2020 SARS-COV-2 seed grant to R.E.A and NIAID grant R01-AI127521 awarded to J.S.M. We are grateful for the efforts of the Texas Advanced Computing Center (TACC) Frontera team and for the compute time made available through a Director’s Discretionary Allocation (made possible by the National Science Foundation award OAC-1818253). We also thank Dr. Fiona Kearns for assistance with the NTD glycan modeling.

## AUTHOR CONTRIBUTIONS

EA and RR conceived the escape mutant procedure. EA, GP, DL, LC, NVJ, IP, SDM, AM, RM, EC, IH, LB, REA performed experiments, modelling and analyzed data. EA and RR wrote the manuscript. All authors contributed to the final revision of the manuscript. EA, EM, REA, JSM and RR coordinated the project.

## RESOURCE AVAILABILITY

Further information and requests for resources and reagents should be directed to and will be fulfilled by Rino Rappuoli.

## Materials Availability

Reasonable amounts of reagents will be made available by Rino Rappuoli upon request under a Material Transfer Agreement (MTA) for non-commercial usage.

## Supplementary Materials

## Materials and Methods

### Enrollment of SARS-CoV-2 convalescent donors and human sample collection

COVID-19 convalescent plasma were provided by the National Institute for Infectious Diseases, IRCCS – Lazzaro Spallanzani Rome (IT) and Azienda Ospedaliera Universitaria Senese, Siena (IT). Samples were collected from convalescent donors who gave their written consent. The study was approved by local ethics committees (Parere 18_2020 in Rome and Parere 17065 in Siena) and conducted according to good clinical practice in accordance with the declaration of Helsinki (European Council 2001, US Code of Federal Regulations, ICH 1997). This study was unblinded and not randomized.

### ELISA assay with SARS-CoV-2 S-protein prefusion trimer and S1 – S2 subunits

COVID-19 convalescent plasmas were screened by ELISA to profile their binding to the SARS-CoV-2 S-protein, S1 and S2 subunits (*1*). Briefly, 384-well plates were coated with 3 μg/mL of streptavidin diluted in coating buffer (0.05 M carbonate-bicarbonate solution, pH 9.6) and incubated at room temperature (RT) overnight. Plates were then coated with SARS-CoV-2 S-protein, S1 or S2 subunits at 3 μg/mL and incubated for 1h at RT. 50 μL/well of saturation buffer (PBS with 1% BSA) was used to saturate unspecific binding and plates were incubated at 37°C for 1h without CO_2_. Plasma samples were diluted 1:10 in PBS/BSA 1%/Tween20 0.05% in 25 μL/well final volume and incubated for 1h at 37°C without CO_2_. Following, 25 μL/well of alkaline phosphatase-conjugated goat anti-human IgG (Sigma-Aldrich) was used as secondary antibodies. Wells were washed three times between each step with PBS/BSA 1%/Tween20 0.05%. Finally, pNPP (p-nitrophenyl phosphate) (Sigma-Aldrich) was used as soluble substrate to detect the polyclonal response to SARS-CoV-2 S-protein, S1 or S2 subunit and the final reaction was measured using the Varioskan Lux Reader (Thermo Fisher Scientific) at a wavelength of 405 nm. Samples were considered as positive if OD at 405 nm (OD_405_) was twice the blank.

### SARS-CoV-2 authentic virus neutralization assay

The mAbs and plasma neutralization activity was evaluated using a CPE-based assay as previously described (*2*). mAbs were tested at a starting concentration of 1 μg/mL diluted in steps of 1:2 while plasma samples were tested at a starting dilution of 1:10 and then diluted in steps of 1:2 for twelve points. All samples were mixed with a SARS-CoV-2 WT, SARS-CoV-2 D614G or SARS-CoV-2 PT188-EM viral solution containing 100 TCID_50_ of the virus. After 1 hour incubation at 37°C, 5% CO_2_, virus-mAb mixture was added to the wells of a 96-well plate containing a sub-confluent Vero E6 cell monolayer. Plates were incubated for 3 days at 37°C in a humidified environment with 5% CO_2_, then examined for CPE by means of an inverted optical microscope. Absence or presence of CPE was defined by comparison of each well with the positive control (plasma sample showing high neutralizing activity of SARS-CoV-2 in infected Vero E6 cells) and negative control (human serum sample negative for SARS-CoV-2 in ELISA and neutralization assays and Vero E6 alone). Technical triplicates were performed for each experiment.

### Cell culture conditions

African green monkey kidney Vero E6 cells (American Type Culture Collection [ATCC] #CRL-1586) were grown in Dulbecco’s Modified Eagle’s Medium (DMEM) high glucose supplemented with 2 mM L-glutamine, 100 U/mL of penicillin, 100 μg/mL streptomycin (“complete DMEM” medium) and 10% fetal bovine serum (FBS). Cells were cultured at 37°C, 5% CO_2_ and passaged every 3-4 days. 18-24 hours before execution of the viral escape assay, plates and propagation flasks containing a standard concentration of Vero E6 cells were prepared in complete DMEM medium supplemented with 2% FBS and incubated at 37°C, 5% CO_2_ until use. 24-well plates were inoculated with 2×10^4^ cells/well to passage the virus-antibody mixture and a virus-only control. 25 cm^2^ flasks pre-seeded with 1×10^5^ cells/mL were prepared in parallel to propagate the viral strains of each experiment to obtain a suitable virus concentration for RNA extraction and subsequent sequencing or RT-PCR analysis. 96 well-plates were inoculated with 1.5×10^4^ cells/well and used for titration of the virus-antibody mixture at each passage.

### Virus propagation and titration

Wild-type SARS CoV-2 2019 (2019-nCoV strain 2019-nCov/Italy-INMI1) virus was purchased from the European Virus Archive goes Global (EVAg, Spallanzani Institute, Rome). For virus propagation, 175 cm^2^ flasks were seeded with Vero E6 cells diluted in complete DMEM high glucose supplemented with 2% FBS at a concentration of 1×10^6^ cells/mL, and incubated at 37°C, 5% CO_2_ for 18-20 hours. After 2 washes with sterile Dulbecco’s phosphate-buffered saline (DPBS), the sub-confluent cell monolayer was inoculated with the SARS-CoV-2 virus at a multiplicity of infection (MOI) of 0.001, incubated for 1 hour at 37°C, 5% CO_2_. Flasks were then filled with 50 mL of complete DMEM 2% FBS, incubated at 37°C, 5% CO_2_, and checked daily until approximately 80-90% of the cell culture showed cytopathic effect (CPE). Supernatants of the infected culture were collected and centrifuged at 4°C, 1600 rpm for 8 minutes to remove cell debris, aliquoted and stored at −80°C. A titration of the propagated viral stocks was performed in 96-well plates containing confluent Vero E6 monolayers, using a 50% tissue culture infectious dose assay (TCID_50_). Cells infected with serial 10-fold dilutions (10^−1^ to 10^−11^) of the virus were incubated at 37°C, 5% CO_2_ and monitored for signs of virus-induced CPE under an inverted optical microscope for 3–4 days. The viral titer, defined as the reciprocal of the highest viral dilution resulting in at least 50% CPE in the inoculated wells, was calculated with the Spearman-Karber formula (*3*).

### Viral escape assay using authentic SARS-CoV-2

To detect neutralization-resistant SARS-CoV-2 escape variants, a standard concentration of the virus was sequentially passaged in cell cultures in the presence of serially diluted samples containing SARS-CoV-2-specific antibodies. Briefly, 12 serial 2-fold dilutions of PT188 plasma prepared in complete DMEM 2% FBS (starting dilution 1/10) were added to the wells of one 24-well plate (final volume in well: 550 μL). 550 μL of virus solution containing 10^5^ TCID_50_ of authentic SARS-CoV-2 was dispensed in each antibody-containing well and the plates were incubated for 1 hour at 37°C, 5% CO_2_. The mixture was then added to the wells of a 24-well plate containing a sub-confluent Vero E6 cell monolayer. Plates were incubated for 5–7 days at 37°C, 5% CO_2_ and examined for the presence of CPE using an inverted optical microscope. A virus-only control and a cell-only control were included in each plate to assist in distinguishing absence or presence of CPE. At each virus passage, the content of the well corresponding to the lowest sample dilution that showed complete CPE was diluted 1:10 and 110 μL/well of this solution was transferred to the antibody-containing wells of the pre-dilution 24-well plate prepared for the subsequent virus passage. After 1 hour incubation at 37°C, 5% CO_2_, the virus-antibody mixture was transferred to Vero E6-containing plates and incubated as before. The virus pressured with PT188 was passaged in cell cultures along with freshly prepared PT188-solution until manifestation of CPE at lower antibody dilutions. At each passage, both the virus pressured with PT188 and the virus-only control were harvested, propagated in 25cm^2^ flasks and aliquoted at −80°C to be used for RNA extraction, RT-PCR and sequencing. The virus-only control was passaged in parallel and in the absence of antibody to help distinguish between adaptation to cell culture conditions and presence of escape mutations. Parallel titrations of the antibody-pressured virus was performed in 96-well plates containing sub-confluent Vero E6 cells as previously described to monitor the viral titer at each passage.

### RNA Extraction

To isolate the viral genetic material for NGS and detection by RT-qPCR, an RNA extraction step was performed using CommaXP® virus DNA/RNA extraction (Spin Column) commercial kit, according to manufacturer’s instruction. Briefly, 500 μL of “Buffer GLX” was added to 300 μL of sample, vortexed for 1 minute and incubated at RT for 5 minutes to allow virus inactivation. The mixture was then added into a spin column inserted in a collection tube and centrifuged at 12,000 rpm for 1 minute at RT. The eluted solution was discarded and 500 μL of “Buffer PD” previously re-suspended in isopropanol were added to the column, then centrifuged as before. Following elimination of the eluted solution, the column was washed with 700 μL of “Buffer PW” previously re-suspended in absolute ethanol and centrifuged as before. This wash was repeated twice. The spin column was then centrifuged at 12,000 rpm for 2 minutes and left with open lid for 5 minutes to allow evaporation of residual ethanol. The column was placed in a new collection tube and 100 μL of RNAse-free ddH_2_0 were added. After a 2 minute incubation at RT, the column was centrifuged for 2 minutes at 12,000 rpm to elute the RNA, which was collected in a new tube for PCR analysis.

### Library preparation and sequencing

Each viral RNA was retrotranscribed using Superscript IV First-Strand Synthesis System (Thermo Fisher Scientific) without the optional RNase H step and used as input for sequencing library construction. Library preparation was performed with the swift amplicon SARS-CoV-2 research panel (Swift Biosciences, Ann Arbor, MI USA) according to manufacturer’s instructions. Library preparation workflow requires two sequential PCR steps. First, more than 341 specific regions are selectively amplified in a targeted single multiplex PCR amplification reaction. Next, a universal PCR is performed to label all amplicons with unique combinations of dual indexed adapters, enabling multiplexing of samples in the same run. Bead-based clean-ups were used to purify the sample by removing unused oligonucleotides and changing buffer composition between steps. Purified individually tagged libraries were quantified by qPCR using Kapa Lib Quant Kit (Roche Diagnostics). In conjunction with the qPCR Ct values we used a library size of 265 bp to calculate library molarity. All the obtained libraries passed quality check and were quantified before being pooled at equimolar concentration and sequenced on Illumina MiSEQ 2×250bp paired-end mode following standard procedures including “Adapter Trimming” and “Adapter Trimming Read 2” option. Sequenced reads were quality trimmed using Trimmomatic software during data analysis. Only good quality reads were mapped against SARS-CoV-2_human_ITA_INMI1_2020 GenBank: MT066156.1 using BWA software with default parameters. After inspection using IGV software, consensus sequences were created for each processed sample.

### RT-PCR

Multiplex real time RT-PCR for simultaneous detection of SARS-CoV-2-2019 N gene and RdRp gene was performed using NeoplexTM COVID-19 Detection Kit. The primer and probe system of the kit is based on the standard TaqMan® Technology. SARS-CoV-2 specific probes are labelled with the FAM and JOE fluorophore to target COVID-19 RdRp and N genes, respectively. The internal PCR control contains primers for targeting human RNaseP mRNA and probes labelled with the Cy5 fluorophore. For RT-PCR, 5 μL of extracted RNA, 5 μL of DW/RNase-free water, 5 μL of COVID-19 PPM (containing primers and probes for targeting RdRp gene, N gene, and human RNase mRNA as an internal control) and 5 μL of One-step Master Mix were used in a final reaction volume of 20 μL to be run in a LightCycler® 96 System (Roche). A negative control consisting of RNase-free water, and a COVID-19 Positive Control (which includes RdRp, N, and internal control target genes as in vitro transcript (IVT) RNA) were included in each run. The PCR cycling conditions used were as follows: reverse transcription was performed at 53°C for 2 minutes, then an initial amplification was done with a denaturation step at 95°C for 2 min, followed by 40 cycles of denaturation at 95°C for 3 sec and primer annealing/extension at 60°C for 30 s. Reactions were run in duplicate in the same experiment. Data were collected by the LightCycler software during the annealing phase of each cycle of amplification. For each sample, a cycle threshold (Ct) was generated for each target (N gene, RdRp and internal control), based on the cycle number where the instrument software detected a log increase in fluorescence of the given sample.

### Negative stain electron microscopy

SARS-CoV-2 S-protein was expressed and purified as previously described (*4*). Purified spike was combined with individual Fabs at final concentrations of 0.04 mg/mL and 0.16 mg/mL, respectively. Following a 30-minute incubation on ice, each complex was deposited on plasma cleaned CF-400 grids (EMS) and stained using methylamine tungstate (Nanoprobes). Grids were imaged at 92,000X magnification in a Talos F200C TEM microscope equipped with a Ceta 16M detector (Thermo Fisher Scientific). CTF estimation and particle picking were performed using cisTEM (*5*), and particle stacks were exported to cryoSPARC v2 (*6*) for 2D classification, *ab initio* 3D reconstruction, and heterogeneous refinement.

### Computational Methods

The PT188-EM spike escape mutant was modeled using *in-silico* approaches. As the mutations are localized in two different domains of the spike, namely the NTD and the RBD, separate models were generated for each domain. In detail, two models of the PT188-EM spike NTD (residues 13-308) were built starting from two different cryo-EM structures of the wild type S protein as templates: (i) one bearing a completely resolved NTD (PDB ID: 7JJI (*7*)), which includes all the loops from N1 to N5, and (ii) one bound to the antibody 4A8 (PDB ID: 7C2L (*8*)), which presents only one small gap within the N5 loop. The model of the PT188-EM spike RBD (iii) was based on the cryo-EM structure of the spike’s RBD in complex with ACE2 (PDB ID: 6M17 (*9*)). The generated models were subsequently refined using explicitly solvated all-atom molecular dynamics (MD) simulations. The 11-amino-acid insertion between Y248 and L249 within the NTD was modeled as a loop using Modeller9.19 (*10*) and keeping all the original cryo-EM coordinates fixed but allowing a certain extent of flexibility for the flanking residues, namely Y248 and 249-257. The F140 deletion was modeled at the same time as the insertion, only allowing residues 138-141 to be flexible. For both the two PT188-EM spike NTD constructs (i.e., one based on 7JJI and one upon 7C2L), 500 models were independently generated, and the system with the best (i.e., the lowest value) Z-DOPE (Discrete Optimized Protein Energy) score was selected upon visual inspection. The modeled PT188-EM spike NTDs were fully glycosylated at the native N-linked glycosylation sites (N17, N61, N74, N122, N149, N165, N234, N282) and at the new N-linked sequon N248d-K248e-S248f introduced within the 11-amino-acid insertion. The glycosylation profile was chosen to be consistent with available glycoanalytic data (*11, 12*) and analogously to Casalino et al (*13*). Although there is no information available for the new glycan at position N248d, this was modeled as FA2 complex-type, similarly to the glycan at position N149 (FA3 complex-type). Starting conformations for the glycans were derived from Casalino et al. (*13*). For the PT188-EM spike RBD carrying E484K mutation we used a model of the fully glycosylated spike in complex with the human ACE2 receptor previously built by Casalino et al. (*14*) and based on the RBD/ACE2 complex simulated by Barros et al. (*15*), which in turn was modelled upon the cryo-EM structure by Yan et al. (*9*) Using this construct, E484 was mutated into a lysine using PSFGEN within VMD (*16*). The two glycosylated constructs for the PT188-EM spike NTD and the one accounting for the E484K mutation within the RBD described above were embedded into an orthorhombic box of explicit waters with a 150 mM concentration of Na+ and Cl-ions, leading to a final size of (i) _~_185,151, (ii) 190,246 and (iii) ~1,178,601 atoms, respectively. Protonation states were assessed using PROPKA3 (*17*) at pH 7.4. The final set up was done with PSFGEN and VMD (*16*), whereas MD simulations were run on TACC Frontera computer facility using NAMD 2.14 (*18*) and CHARMM36m force fields to refine the models (*19–21*). All the simulations were carried out using a 2 fs timestep with SHAKE (*22*) algorithm to keep the bonds involving hydrogen atoms fixed. The cutoff for non-bonded van der Waals and short-range electrostatic interactions was set to 12 Å, whereas the particle-mesh Ewald (*23*) approach was employed to account for long-range electrostatics. All three systems were first minimized using the conjugate gradient energy approach for 10,000 steps. Subsequently the temperature of the systems was gradually increased to 310 K for 1 ns in NVT ensemble, while imposing a harmonic restraint of 5 kcal/mol to the protein and the glycan atoms. Next, the restraints were released, and the systems were coupled to a Langevin thermostat (*24*) (310 K) and a Nosé-Hoover Langevin piston (*25, 26*) (1.01325 bar) and equilibrated for ~5 ns in NPT conditions. After this point only the PT188-EM spike NTD model (i) based on 7JJI was subjected to production run MD for ~100 ns to check for the flexibility of the 11 aa insertion loop and the impact of the F140 deletion, whereas the simulations of the other PT188-EM spike NTD model (ii) based on 7CL2 and of the PT188-EM spike RBD model were stopped. The systems and the simulations were visually inspected with VMD, which was also used for image rendering (*16*).

## SUPPLEMENTARY TABLES

**Table S1.**
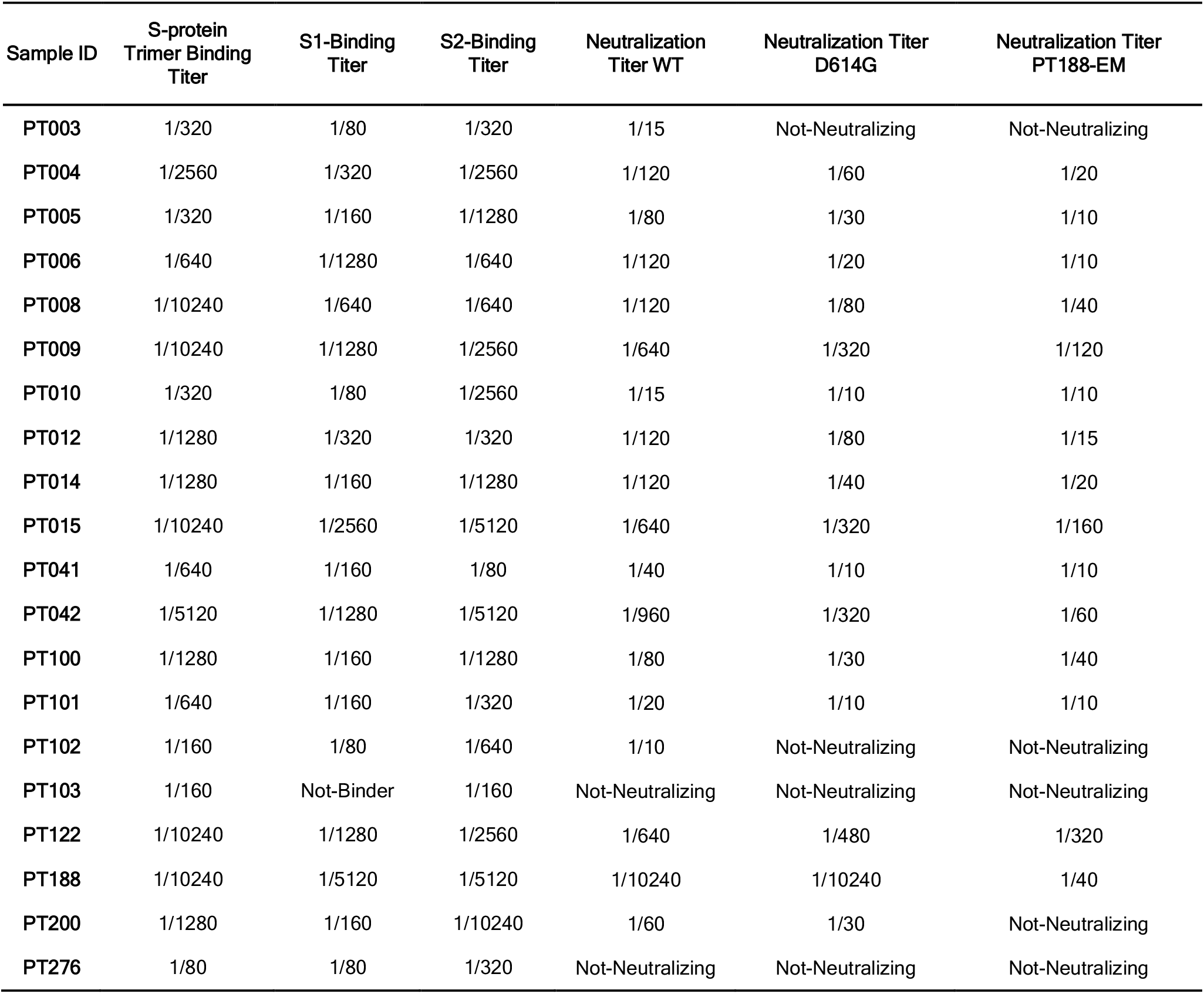
Summary of COVID-19 convalescent plasma characteristics. The table shows the binding profile and neutralization activities of twenty COVID-19 convalescent plasma samples.

**Table S2.** SARS-CoV-2 escape mutant summary. The table shows the starting dilution, passages, viral titer and neutralization activity of the plasma per each passage to generate the authentic virus SARS-CoV-2 escape mutant.

| Sample ID | Starting Dilution | Passage | Days of Incubation<br>(Days per Passage) | Viral Titer | Mutation<br>(% Virion Frequency) | Neutralization<br>Titer |
| --- | --- | --- | --- | --- | --- | --- |
| PT188 | 1/10 | 0 | 5 (5) | $10^{5.4}$ | - | 640 |
| PT188 | 1/10 | 1 | 10 (5) | $10^{5.4}$ | - | 640 |
| PT188 | 1/10 | 2 | 15 (5) | $10^{5.7}$ | - | 640 |
| PT188 | 1/10 | 3 | 20 (5) | $10^{5.7}$ | - | 640 |
| PT188 | 1/10 | 4 | 25 (5) | $10^{5.2}$ | - | 640 |
| PT188 | 1/10 | 5 | 31 (6) | $10^{5.1}$ | - | 640 |
| PT188 | 1/10 | 6 | 38 (7) | $10^{4.9}$ | - | 640 |
| PT188 | 1/10 | 7 | 45 (7) | $10^{4.2}$ | Deletion F140 - NTD (39%) | 320 |
| PT188 | 1/10 | 8 | 53 (8) | $10^{4.3}$ | Deletion F140 - NTD (100%) | 160 |
| PT188 | 1/10 | 9 | 59 (6) | $10^{4.4}$ | - | 160 |
| PT188 | 1/10 | 10 | 66 (7) | $10^{4.5}$ | - | 160 |
| PT188 | 1/10 | 11 | 73 (7) | $10^{4.0}$ | - | 160 |
| PT188 | 1/10 | 12 | 80 (7) | $10^{3.9}$ | Substitution E484K - RBD<br>(100%) | 40 |
| PT188 | 1/10 | 13 | 85 (5) | $10^{5.7}$ | Insertion<br>248aKTRNKSTSRRE<sub>248k</sub> - NTD<br>(49%) | Not-neutralizing |
| PT188 | 1/10 | 14 | 93 (8) | $10^{5.4}$ | Insertion<br>248aKTRNKSTSRRE<sub>248k</sub> - NTD<br>(100%) | Not-neutralizing |

**Table S3.** Features of thirteen SARS-CoV-2 neutralizing antibodies. The table shows the binding and neutralization profile of thirteen previously identified SARS-CoV-2 nAbs. Asterisked columns refer to previously published data (*1*).

| mAb ID | Binding specificity* | Neutralization WT IC<sub>100</sub> (ng:mL)* | Neutralization D614G IC<sub>100</sub> (ng:mL)* | Neutralization PT188-EM IC<sub>100</sub> (ng:mL) |
| --- | --- | --- | --- | --- |
| J08 | S1-RBD | 3.9 | 7.8 | 22.1 |
| I14 | S1-RBD | 11.0 | 19.7 | 176.8 |
| F05 | S1-RBD | 3.9 | 4.9 | Not-Neutralizing |
| G12 | S1-RBD | 39.4 | 39.4 | Not-Neutralizing |
| C14 | S1-RBD | 157.5 | 78.7 | 88.4 |
| B07 | S1-RBD | 99.2 | 49.6 | 250.0 |
| I21 | S1 | 99.2 | 198.4 | Not-Neutralizing |
| J13 | S1 | 396.8 | 500.0 | Not-Neutralizing |
| D14 | S1 | 396.8 | 250.0 | Not-Neutralizing |
| H20 | S-protein | 492.2 | 310.0 | Not-Neutralizing |
| I15 | S-protein | 310.0 | 155.0 | Not-Neutralizing |
| F10 | S-protein | 155.0 | 195.3 | Not-Neutralizing |
| F20 | S-protein | 246.1 | 155.0 | Not-Neutralizing |

## SUPPLEMENTARY FIGURES

**Figure S1.**
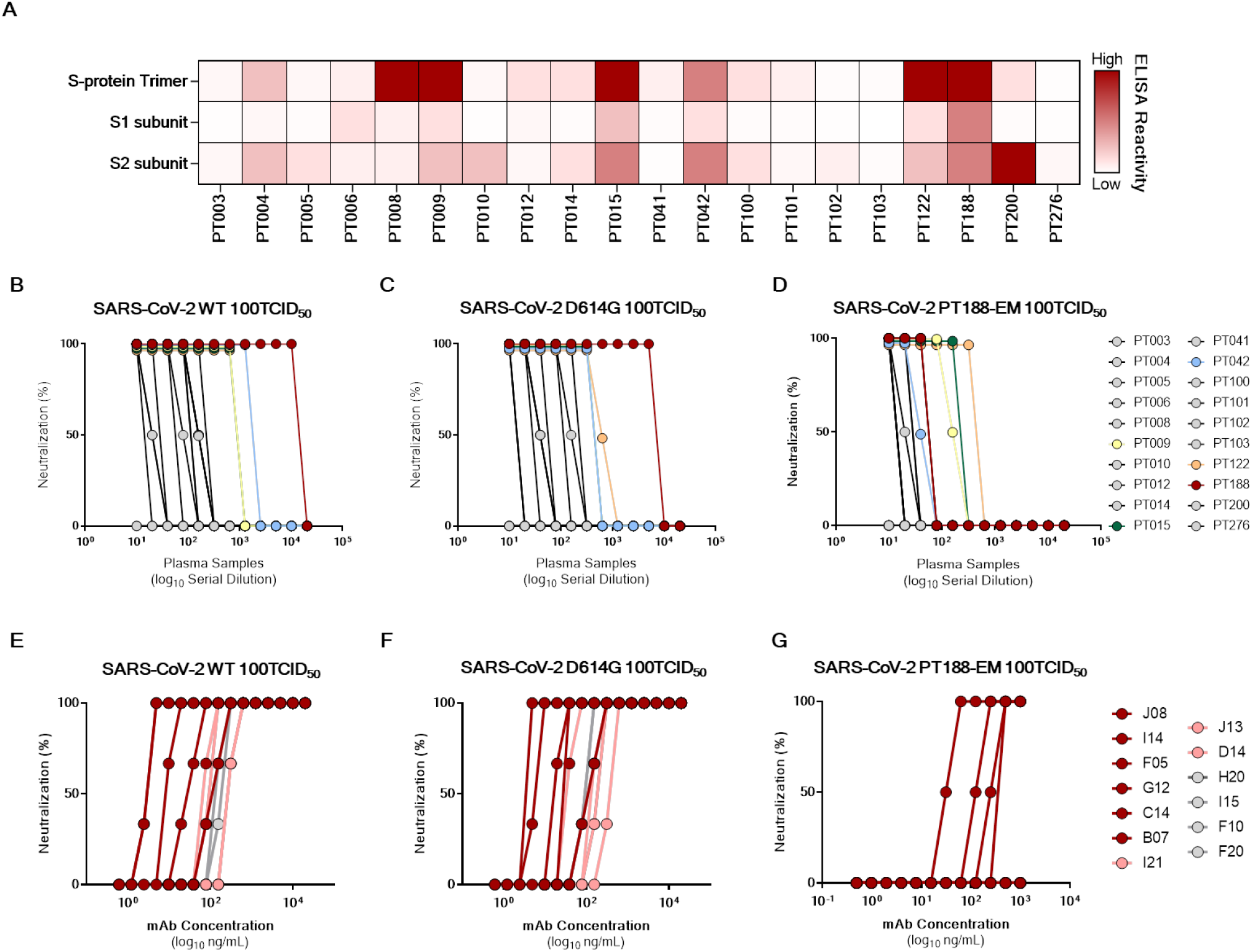
Binding and neutralization profiling of plasma samples from twenty COVID-19 convalescent patients. (A) Binding profiling of COVID-19 convalescent plasma to S-protein trimer, S1 subunit and S2 subunit. (B – D) Neutralization activity of COVID-19 convalescent plasma against SARS-CoV-2 WT (B) and D614G (C) viruses and SARS-CoV-2 PT188-EM (D). Data are representative of technical triplicates. All plasma are shown in grey, only most potently neutralizing plasma were colored in yellow, green, light blue, orange and red for PT009, PT015, PT042, PT122 and PT188 respectively. (E – G) Neutralization curves for the thirteen tested mAbs against SARS-CoV-2 WT (E), SARS-CoV-2 D614G (F) and SARS-CoV-2 PT188-EM respectively (G). Data are representative of technical triplicates.

**Figure S2.**
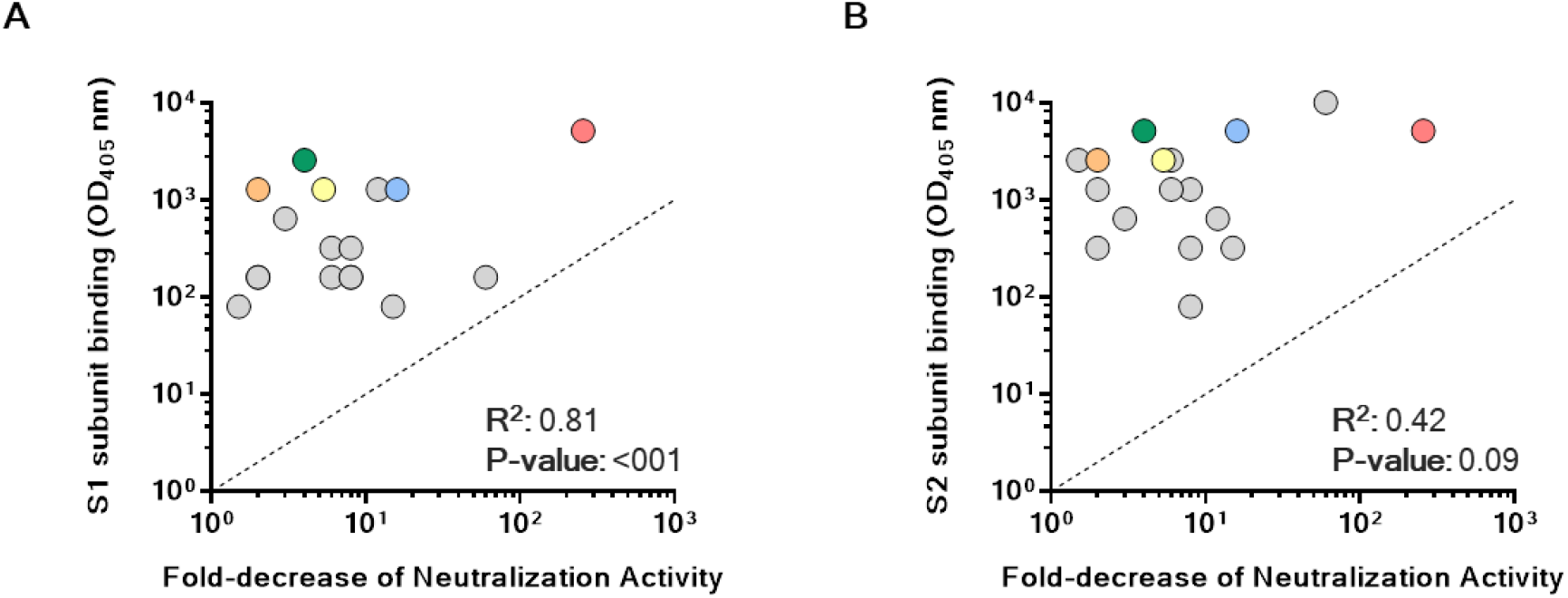
Binding and functional characterization of COVID-19 convalescent plasma. (A – B) Graphs show correlation between S1 subunit (A) and S2 subunit (B) binding and fold-decrease of neutralization activity. All plasma are shown in grey, only most potently neutralizing plasma were colored in yellow, green, light blue, orange and red for PT009, PT015, PT042, PT122 and PT188 respectively.

**Figure S3.**
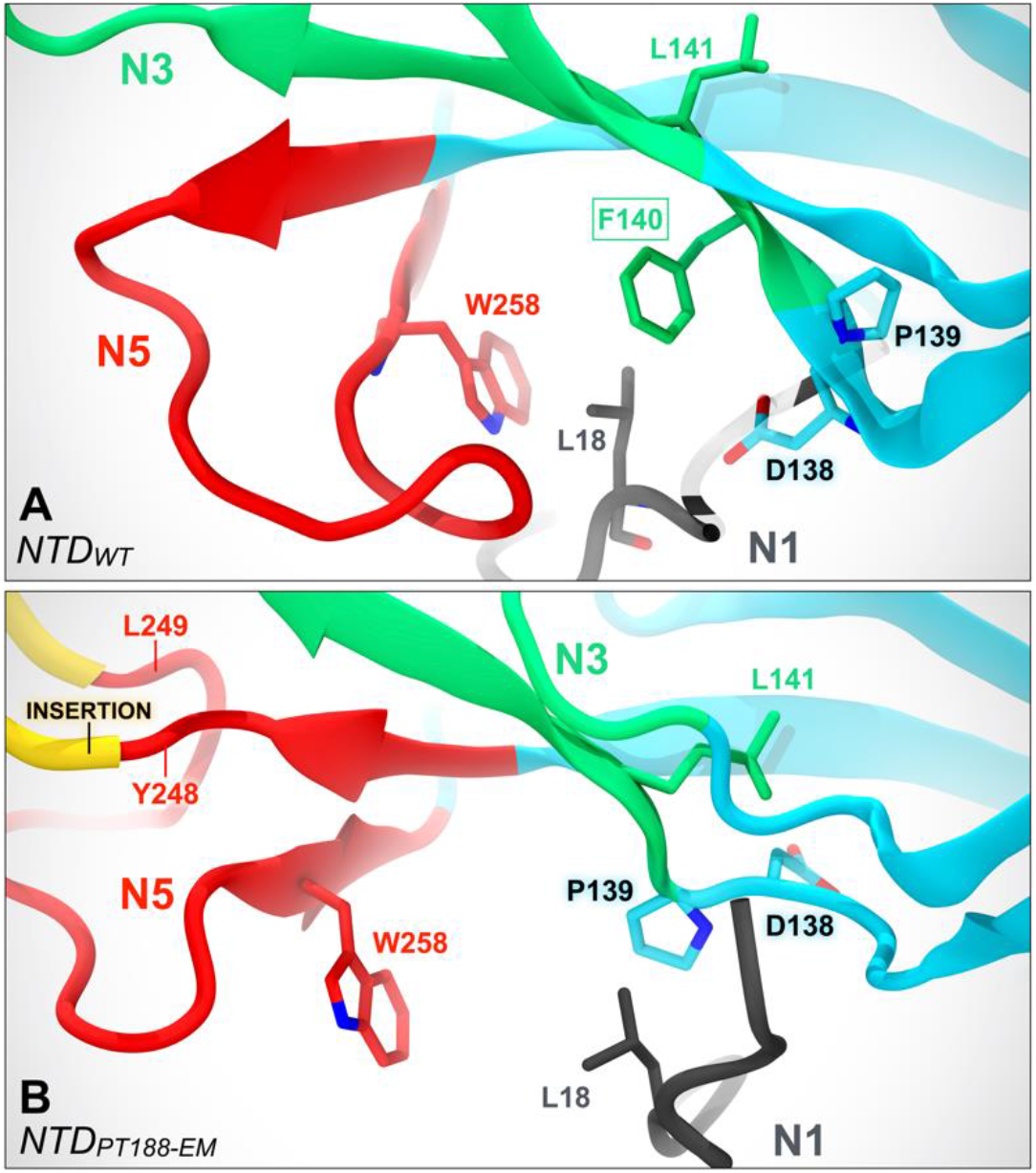
F140 deletion affects packing of N1, N3 and N5 loops. (A) Molecular representation of the NTD of the WT SARS-CoV-2 S protein as in the cryo-EM structure with PDB id 7JJI. F140 (N3) establishes hydrophobic contacts with L18 (N1) and W258 (N5). (B) Molecular representation of the model of the PT188-EM spike NTD after molecular dynamics simulations. The pattern of interactions is disrupted by the deletion of F140, leading to a loosening of the N1/N3/N5 loop packing.

**Figure S4.**
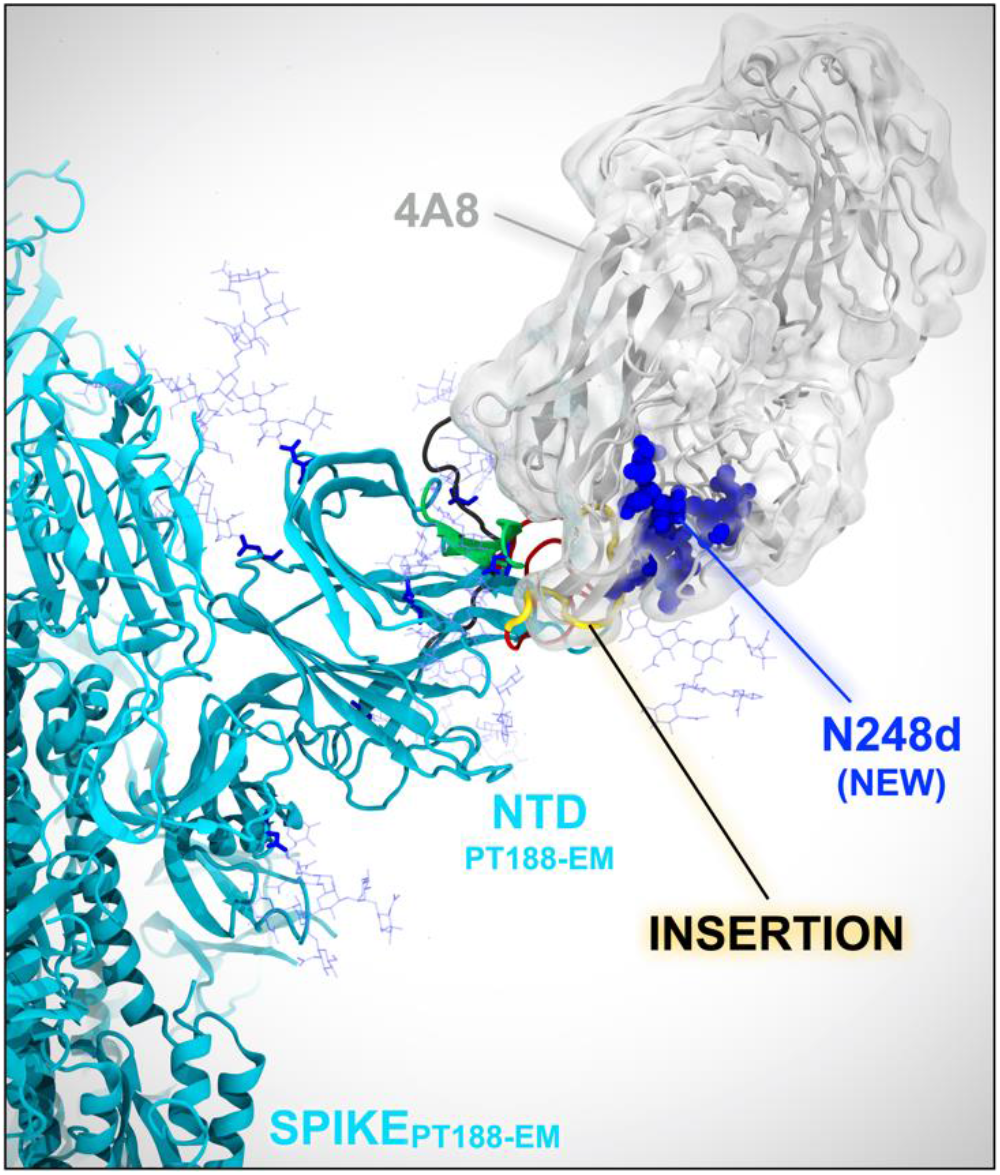
Insertion of 11 amino acids introduces a new N-glycan within N5 loop. (A) Molecular representation of the molecular dynamics-equilibrated model of the PT188-EM spike NTD based on PDB id 7C2L. The original cryo-EM structure used for this model already provided the coordinates for the NTD-bound 4A8 antibody (gray transparent surface). The relaxed model was aligned onto the cryo-EM coordinates, therefore retrieving the initial 4A8-bound pose, allowing for evaluation of steric compatibility.

**Figure S5.**
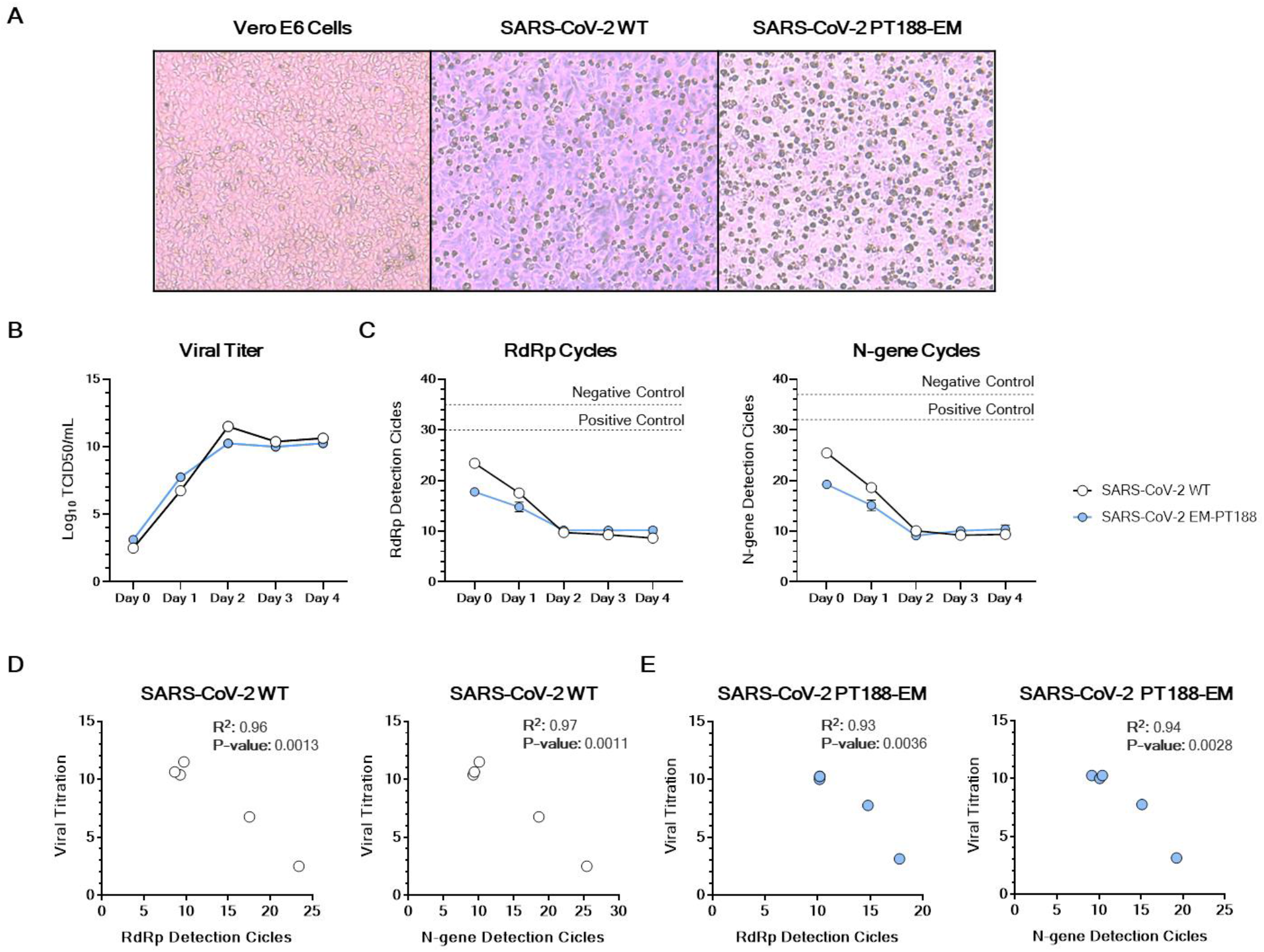
Evaluation of SARS-CoV-2 PT188-EM viral fitness. (A) Representative microscope images showing Vero E6 feeder cells alone and CPE observed for the SARS-CoV-2 WT and SARS-CoV-2 PT188-EM. (B) Viral titer of SARS-CoV-2 WT (white dots) and SARS-CoV-2 J08-EM (red dots) viruses shown as Log_10_ TCID_50_/mL. (C) RT-PCR detection cycles for RdRp and N-gene. Dotted lines represent the negative and positive control thresholds per each gene. Curves show technical duplicates. (D – E) Correlation between viral titer and RdRp or N-gene detection cycles for SARS-CoV-2 WT (white dots) and SARS-CoV-2 PT188-EM (red dots) respectively.

